# Multi-site clonality analyses uncovers pervasive subclonal heterogeneity and branching evolution across melanoma metastases

**DOI:** 10.1101/848390

**Authors:** Roy Rabbie, Naser Ansari-Pour, Oliver Cast, Doreen Lau, Francis Scott, Sarah J. Welsh, Christine Parkinson, Leila Khoja, Luiza Moore, Mark Tullett, Kim Wong, Ingrid Ferreira, Julia M. Martínez Gómez, Mitchell Levesque, Ferdia A. Gallagher, Alejandro Jiménez-Sánchez, Laura Riva, Martin L. Miller, Kieren Allinson, Peter J. Campbell, Pippa Corrie, David C. Wedge, David J. Adams

**Author notes:** These authors contributed equally to this work (Roy Rabbie, Naser Ansari-Pour). **Correspondence to:** Dr. David Wedge, Big Data Institute, Nuffield Department of Medicine, University of Oxford, Oxford, UK, *, ***Email:***, Dr. David Adams, Experimental Cancer Genetics, Wellcome Sanger Institute, Hinxton, Cambridge, CB10 1HH, *, ***Email:***. **Author conflicts of interests:** No competing interests declared.

## Abstract

Metastatic melanoma carries a poor prognosis despite modern systemic therapies. Understanding the evolution of the disease could help inform patient management. Through whole-genome sequencing of 13 melanoma metastases sampled at autopsy from a treatment naïve patient and by leveraging the analytical power of multi-sample analyses, we reveal that metastatic cells may depart the primary tumour very early in the disease course and follow a branched pattern of evolution. Truncal UV-induced mutations that often swamp downstream analyses of heterogeneity, were found to be replaced by APOBEC-associated mutations in the branches of the evolutionary tree. Multi-sample analyses from a further 7 patients confirmed that branched evolution was pervasive, representing an important mode of melanoma dissemination. Our analyses illustrate that combining cancer cell fraction estimates across multiple metastases provides higher resolution phylogenetic reconstructions relative to single sample analyses and highlights the limitations of accurately inferring inter-tumoural heterogeneity from a single biopsy.

## Background

Large-scale sequencing studies in cutaneous melanoma have revealed the complex mutational landscape of the disease^1–4^. However, few studies have explored the temporal and spatial evolution of molecular alterations acquired during disease progression. Such findings may inform risk prediction and our understanding of the mode of metastatic spread, with implications for future patient management. Current methods for reconstructing clonal evolution from bulk sequencing data rely on computational approaches to identify sets of mutations that have a similar clonal frequency which are assigned to respective clones within the tumour^5^. Recent studies have used these algorithms to infer the evolutionary relationship between clones across multiple samples, correlating these insights with changes in disease progression or therapy response. The fraction of cancer cells carrying each cluster of mutations in each tumour (cancer cell fraction, CCF) represents the frequency of the corresponding subclones in that sample. By comparing the constituent subclonal mutations between pairs of tumours, it is possible to derive the ancestral relationships between subclones rather than between samples, thereby constructing true phylogenies^6^. This type of quantitative modelling provides much greater resolution than single-sample studies and has yielded important insights into the patterns and timing of tumour cell spread^7–11^. In particular, two subclones can be either linearly related to each other, or have a common ancestor but develop on opposing branched lineages (herein referred to as branched evolution). We note that throughout this study, we refer to mutations observed in all tumour cells within a sample as ‘clonal’, those found in a subset of tumour cells as ‘subclonal’ and those found clonally in all samples from the same patient as ‘truncal’.

The mutation rate of melanoma is one of the highest among all malignancies^12^ and, as somatic mutations mirror a cancer’s initiation and evolution, genome sequencing studies can provide valuable insights into the progression of the disease. Using targeted panel sequencing of 263 cancer driver mutations across 12 primary melanomas matched with regional metastases, Shain and colleagues^13^ demonstrated that whilst some primary melanomas and matching regional metastases have pathogenic mutations in just one branch of the phylogenetic tree, there were no driver mutations exclusive to metastases (i.e. not shared with the primaries)^13^. By further showing that most somatic alterations (point mutations and copy number changes) were shared, the authors concluded that primary melanomas and melanoma metastases tend to select for the same set of pathogenic mutations. One feature of such studies is that the clonal composition of each sample is determined using just the presence or absence of mutations in each sample. However, this type of modelling relies on the estimation of clonal frequencies, which is vital for the identification of more than two clones per sample and for accurate phylogenetic reconstructions^14^. A recent whole-exome sequencing (WES) study of 86 distant metastases obtained from 53 patients used the relative variant allele frequency (VAF, frequency of a mutation in sequencing data) of shared versus private mutations in each lesion to infer the likely clonal status of private mutations within each sample^15^, finding that although many private mutations were subclonal, polyclonal seeding (defined as a sample harbouring subclonal mutations represented across distinct clonal lineages) was a rare event^15^. A picture has therefore emerged whereby the majority of mutations in melanoma metastases are truncal and shared by all progeny. Leading up to the formation of a primary melanoma a stepwise model of linear development has been proposed, which includes selection for particular advantageous molecular alterations (including copy number aberrations), facilitating the sequential transition through successive stages^16,17^. Although this is a well-established model in the progression from pre-malignant melanocytic precursors to invasive primary melanomas^17,18^, the evidence for its ubiquity in metastatic progression is less conclusive.

Multi-site sequencing studies in melanoma have thus far been based on a small number of single nucleotide variants (SNVs) falling in coding exons, with gene panels focussed on SNVs in known cancer genes^13,15, 19–23^. Critically, heterogeneity analyses are highly influenced by sequencing coverage and depth, and the limits of detection of mutant alleles. Less precise methods might not enable the detection of the whole catalogue of mutations, particularly heterogenous mutations present in a small percentage of tumour cells, which could lead to an underestimation of tumour heterogeneity. The VAF can also be affected by contributions from alleles in stroma and infiltrating immune cells as well as the presence of both the mutated and wildtype alleles in the tumour. Importantly, changes to the copy number of a locus may also alter the VAF dramatically and, if not accounted for, will result in inaccurate clonal frequency estimates, giving a misleading picture of the clonal structure of a tumour^14^. For example, a mutation that has occurred before a copy number gain is carried by two out of three chromosomal copies, whereas a mutation that occurred after the gain is carried by one out of three copies. Indeed, whole-genome duplication and other copy number aberrations vary across melanoma metastases from the same patient, evolutionary changes that may not be evident from the analysis of SNVs alone^15^. Inferring clonality from allelic frequencies therefore requires an integrative approach harnessing the most sensitive sequencing technologies, while considering measures of tumour ploidy and purity.

In this study, we present the first genome-wide analysis of multiple melanoma metastases sampled at autopsy from a treatment-naïve patient. Using multi-sample clonality analyses in this whole-genome sequenced patient, as well as a further 7 whole-exome sequenced patients, we identify clusters of co-occurring truncal, clonal and subclonal mutations across multiple samples, and uncover the chronological sequence of genomic alterations. We show that metastases in different organs have distinct clonal lineages and conclude that clonal heterogeneity and branched evolution likely predominate in melanoma metastases.

## Results

### An aggressive clinical course characterised by rapidly progressive multi-organ metastases

Our index case was a 71y old male of European descent with no relevant family history, who initially presented with a 1.2mm Breslow thickness non-ulcerated, Clark level 3 superficial spreading melanoma which was resected from the anterior chest wall with a wide local excision (**Fig. 1A**). The patient declined a sentinel lymph node biopsy and staging scans were clear for distant metastases. Five years later, the patient presented to the emergency department with sudden onset receptive dysphasia and dyspraxia. A contrast computerised tomography (CT) head scan showed multiple enhancing lesions in both cerebral hemispheres with adjacent vasogenic oedema consistent with metastases (**Fig. 1B**). A staging contrast CT also showed multiple lung, liver and retroperitoneal lymph node metastases (**Fig. 1B**). A biopsy of one of the liver lesions confirmed metastatic melanoma. Mutation-specific immunohistochemistry for *BRAF^V600E^* did not detect the mutated protein, but targeted panel sequencing identified an activating *BRAF*^V600R^ mutation. However, in view of poor overall performance status and on discussion with the patient and his family, he chose to be managed with best supportive care. The patient consented to undergo a research autopsy as part of the ethically-approved MelResist study (see **Methods**). He received corticosteroids with marked improvement in neurological symptoms and underwent whole-brain radiotherapy 30Gy in 10 fractions. A repeat staging CT scan 2 weeks after completing radiotherapy revealed stable brain metastases but widespread progression of the extracranial disease (**Fig. 1B**). He died four weeks later.

**Fig. 1.**
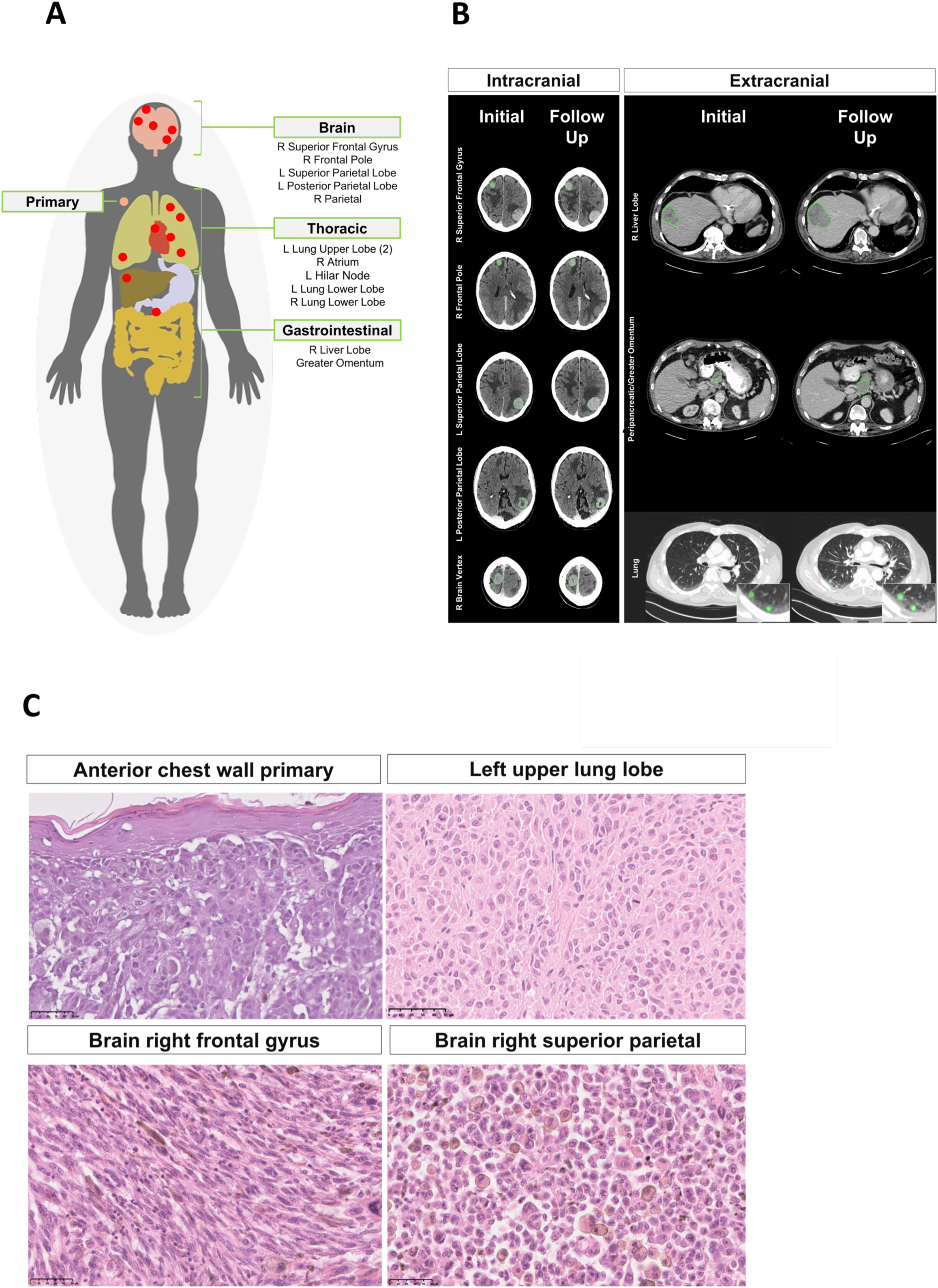
Clinical presentation and sequalae of index autopsy case. A) Sites of 13 metastases sampled during the autopsy and original anterior chest wall cutaneous primary melanoma. B) Axial CT imaging from the brain, chest and abdomen before (left) and after (right) whole brain radiotherapy (imaging 5 weeks apart). Brain CT images represent the following metastatic sites (from top to bottom); right superior frontal gyrus, right frontal pole, left superior parietal, left posterior parietal and right brain vertex (corresponding to the ‘right parietal’ sample labelled in the remainder of the text). Chest/abdomen CT images represent the following metastatic sites (from top to bottom); right lobe of liver, peripancreatic/greater omentum (corresponding to the ‘greater omentum’ sample labelled in the remainder of the text) and right lower lung. C) Histological analyses (Hematoxylin-eosin images in 40-fold magnification), from top left to bottom right (D1-D4); primary melanoma from the anterior chest wall (D1), distant metastasis from the left upper lobe of the lung (D2), right frontal gyrus of the brain (D3) and right superior parietal lobule of the brain (D4). Morphological appearances differ across the tumours, including varying cellular morphology and pigmentation. 50um scale bar shown.

During a research autopsy, metastases were identified macroscopically in the brain, lung, liver and retroperitoneum as well as the right atrium, the latter was identified as an 18mm polypoid lesion arising from the endocardial surface (**Fig. 1C**). The cause of death was identified as a saddle pulmonary embolus. In total, 13 metastases were sampled at autopsy and a further 2 samples were obtained from the archived anterior chest wall primary melanoma for further molecular analyses (**Fig. 1 & Supplementary Table**). Histopathological analyses of the metastases confirmed metastatic melanoma. Morphological heterogeneity was observed between metastases based on cellular features (ranging from epithelioid to spindle cell), as well as in the degrees of pigmentation and necrosis (**Fig. 1C**).

### Metastatic melanomas are dominated by UV-induced clonal mutations that dominate phylogenetic reconstructions

Whole genome sequencing of the 13 metastatic tumours sampled at autopsy, which were sequenced to an average depth of 38x, revealed a union list of ∼118,000 SNVs. We detected 1993 putative somatic indels, of which 10 were frameshifts and common to all metastases (**Supplementary Data**). All 13 metastases carried an activating missense *BRAF*^V600R^ mutation (c.1798_1799delGTinAG), as well as mutations in the melanoma driver genes *PTEN^A43T^* and *MAP2K1^G128S^* (the latter is a novel variant not previously reported in the COSMIC database^24/24^), and an invariant splice-site in *ARID2*, all of which were clonal across all metastases. We further explored clonal architecture using the Cancer Cell Fraction (CCF), determined by adjusting the variant allele frequencies of SNVs for copy number aberration (CNA) status and the extent of normal cell contamination (i.e. purity) as previously described^5^. Briefly, multidimensional Bayesian Dirichlet Process-based mutation clustering (ndDPClust^5,25^) was used to identify truncal, clonal and subclonal mutation clusters based on the CCF of the union list of somatic SNVs across all 13 metastases. Using this approach, we found that >90% of all somatic variants were truncal, with only one additional cluster which represented at least 1% of the SNVs (1651, 1.35%), while the other 179 smaller clusters had a mean of only ∼53.13 SNVs. The large truncal cluster was dominated by C>T transitions at dipyrmidines (characteristic of UV-induced mutational damage^26/26^) and was shared across all metastases, implying that clonal heterogeneity was absent and that only one disseminated clone initiated metastatic outgrowth (**Fig. 2A**). We next filtered for artefactual clusters and SNVs within regions of ambiguous copy number status across all samples (see **Methods**) and subtracted variants assigned to the major truncal cluster which uncovered a union list of 2247 unique non-truncal variants (**Fig. 2B**). Overall, 22/2247 of these variants fell within the protein-coding region of the genome of which 14 were protein-altering (all missense). None of these variants were in established cancer driver genes^24^. We undertook a validation experiment by custom capture pull-down sequencing of selected non-silent truncal SNVs present across all 13 metastases (selected as either cancer driver or loss-of-function SNVs, N=652), as well as all 2247 non-truncal SNVs. In this way 99% of truncal SNVs and 92% of all the non-truncal SNVs (called in the original WGS experiment) were observed to be true variants (see **Methods**).

**Fig. 2.**
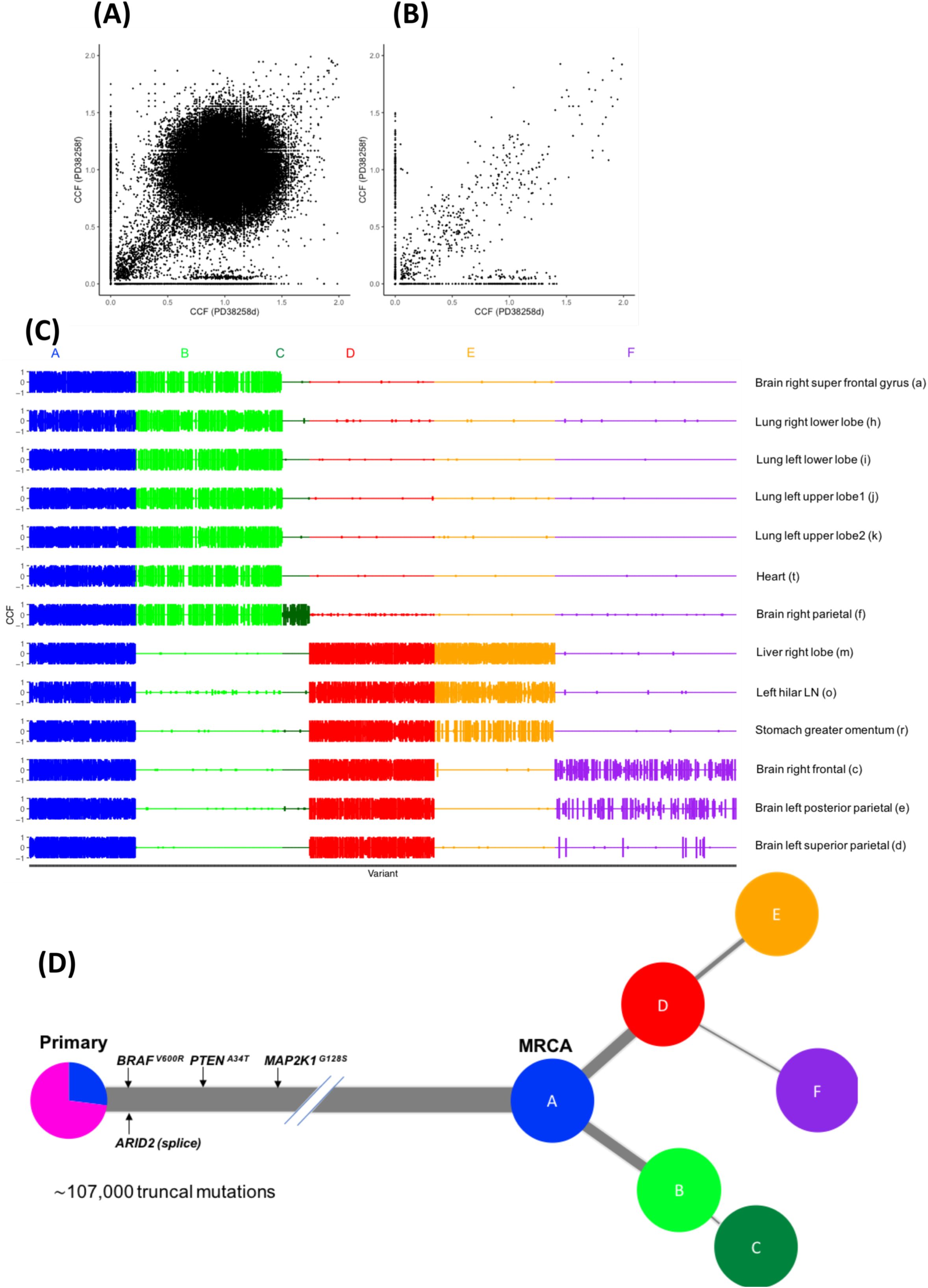
Subtracting the clonal cluster of variants revealed clonal diversification and branching evolution in the index autopsy case. A) Using copy number, tumour purity and ploidy, the variant allele frequency (VAF) of each mutation is converted to the fraction of tumour cells harbouring the mutation, also known as the Cancer Cell Fraction (CCF). Point estimates of the CCF are represented here as black dots across two representative samples in a density plot (left super parietal (PD38258d) and right parietal brain metastases (PD38258f), represented across the X and Y axes respectively). We observed a large cluster of mutations at (1,1), corresponding to SNVs present in all the cells in both sites (CCF=1), indicating truncal variants. B) After removing the truncal variants, the non-truncal mutation clusters uncovered subclonal mutations. Bayesian Dirichlet Process Modelling could then be used to assign mutations to clusters. These clusters help define the distinct populations of cells that arose from clonal expansions during tumour evolution. Although there are still a small number of SNVs at CCF 1, these did not belong to the truncal cluster of mutations, representing those mutations found clonally in all samples. C) The clusters of non-truncal mutations are represented in a CCF distribution plot, wherein rows reflect samples and columns reflect alphabetically and colour-assigned mutation clusters (n=6). The x-axis represents the SNVs within each cluster and the y-axes represent CCF; the latter been plotted in positive and negative directions for improved visualisation of clusters. D) Alphabetically and colour-assigned mutation clusters from (C) are represented in circles, where branch length is proportionate to the number of SNVs in the cluster (filtering SNVs occurring in regions with different CNA status across samples reduced 118,000 SNVs to 107,000 truncal SNVs indicated here) and branch thickness is proportionate to the average CCF of the cluster (across all the samples). Most recent common ancestor (MRCA) represents the final cluster of SNVs carried by all tumour cells in all metastases. Mutations that have occurred before the MRCA are carried by all tumour cells and define the truncal cluster of mutations. Truncal mutations in the melanoma driver genes *BRAF^V600R^, PTEN^A34T^, MAPK2K1^G128S^* (the latter represents a novel mutation not reported in the COSMIC database^24^) and an *ARID2* invariant splice-site are indicated on the trunk of the tree (the order of the driver mutations displayed on the trunk is arbitrary). We observed evidence of branched evolution emanating from a truncal clone. The primary tumour colour shading represents the mutation clusters detected in the primary (using targeted sequencing), whereby the blue represents the subclonal mutation cluster within the primary (at average CCF of 0.27), which were fixed and become truncal across all metastases.

### Reconstructing the phylogenetic tree based on the non-truncal mutation clusters uncovers branched evolution

Applying ndDPClust to the non-truncal variants of the index autopsy case revealed 6 distinct mutation clusters. By assessing the distribution of these clusters as well as the CCF distribution within each cluster across all metastases, we reconstructed the phylogenetic tree of disease evolution (**Fig. 2C & D**) (see **Methods** for further details). The tree split into two main branches (separated by clusters B and D) with one cluster further separating into two additional clusters, so-called ‘leaves’ or terminal branches of the phylogenetic tree (C, E and F). We next assessed the distribution of mutation clusters per sample in order to reconstruct sample-level phylogenetic trees (**Supplementary Fig. 2**). This sample-level tree represents a subtree of the overall phylogenetic tree (**Fig. 2D**) including just those subclones seen within each sample, however in doing this we were able to segregate the samples based on their respective clonal lineage. Analysis in this way revealed two clear lineages, representing distinct waves of metastatic seeding. Interestingly, the brain metastases were represented on both lineages suggesting there were at least two waves of spread to the central nervous system, whereas the lung metastases were derived from a single lineage (**Supplementary Fig. 2**). In summary, we found that mutation clusters from multiple distinct clones were present across multiple metastatic tumours. This approach has revealed, for the first time, convincing evidence of branching evolution in metastatic melanoma. This finding however could not have been otherwise resolved had it not been for the removal of the dominant cluster of truncal variants, which masked this complex phylogenetic architecture.

We next analysed the representation of clonal and subclonal SNVs within two samples from the original cutaneous primary of the index case resected five years earlier, with the aim of tracing back the ancestral clones. We used targeted panel sequencing with a median coverage of 40x, to ascertain whether selected SNVs identified in the genome-sequenced metastatic tumours were also present in the primary, requiring at least 2 supporting reads reporting the alternative allele to call an SNV in the primary (see **Methods**). We found that 573/652 (88%) of the selected truncal variants identified in the metastases could also be detected in the primary, whereas only 1 of the non-truncal cluster variants identified in the metastases was present in the primary (see **Methods**). By selecting the 144/652 truncal SNVs present in diploid regions across all metastases, we were not only able to estimate purity of the two primary samples based on VAF density of SNVs (see **Methods**), we were also able to run ndDPClust on both primary tumour samples by assuming diploidy in SNVs within the primary tumours. In addition to the main clonal cluster, we further identified that the two primary tumour samples harboured the same subclone represented by 37 SNVs (at CCF 0.25 95% CI 0.22-0.37 and 0.29 95% CI 0.18-0.34 for samples PD38258u and PD38258v, respectively), which were fixed and became truncal across the metastases. No known drivers were uniquely present in this primary tumour subclone, however, we did identify a nonsense variant in *IL1R1*, a gene which is thought to act as a tumour suppressor^27^. Taken together, this indicates that the long trunk of the phylogenetic tree originated from this subclonal cluster within the primary tumour, and that the non-truncal metastatic clones arose *de novo* during metastatic dissemination (**Fig. 2D**). These non-truncal metastatic clones that make up the phylogenetic tree therefore represent changes that were acquired in addition to those that establish the primary tumour.

### Multi-site clonality analyses from a further 7 patients revealed pervasive branched evolution across melanoma metastases

In order to assess whether branched evolution was consistent across further cases, we undertook whole-exome sequencing (WES) of 21 melanoma metastases matched with germline blood samples from an additional 7 patients with metastatic melanoma who had consented to take part in the MelResist study. All samples were obtained from clinically or radiologically progressing disease sites, either at the time of first distant relapse, or following systemic therapy with MAP-kinase directed therapies or immune checkpoint inhibitors (**Supplementary Table**). We identified an average of 598 (range of 108-2088) non-synonymous coding variants (including missense, nonsense and splice-region mutations), and 7 (range 2-15) frameshift variants per patient, both totalled across all samples within each patient (**Supplementary Data)**. Six out of 7 patients had metastases arising from a cutaneous primary and 1 patient (MultiSite_WES_Patient1) had an acral primary melanoma. Their metastases (MultiSite_WES_Patient1), as expected, carried a particularly low number of SNVs (only 298 non-synonymous coding variants totalled across 6 metastases). All 7 patients had metastases harbouring an activating *BRAF^V600E^* driver mutation and, in accordance with previous reports^13,15^, all melanoma drivers were represented on the trunks (rather than the branches) of the phylogenetic trees (except for a novel *TP53^R141C^* mutation in patient MultiSite_WES_Patient1) (**Fig. 3**). We again used ndDPClust to cluster SNVs according to their respective CCFs (see **Methods**). We identified 2-10 distinct clusters per patient with clear evidence of branched evolution across 6/7 patients, evidenced by the presence of mutation clusters in mutually exclusive subsets of samples (**Supplementary Fig. 4**). Reconstructing sample-level phylogenetic trees, we identified distinct clonal lineages within each patient (**Supplementary Fig. 4**), and found evidence for polyclonal seeding in two patients. Given that branched evolution was detected in 6/7 cases (including the acral melanoma patient) based on WES – which has a much lower genomic resolution than WGS – and with as little as two samples per patient in most cases (**Supplementary Table**), provides strong support that this mode of clonal evolution is likely to be pervasive in melanoma dissemination.

**Fig. 3.**
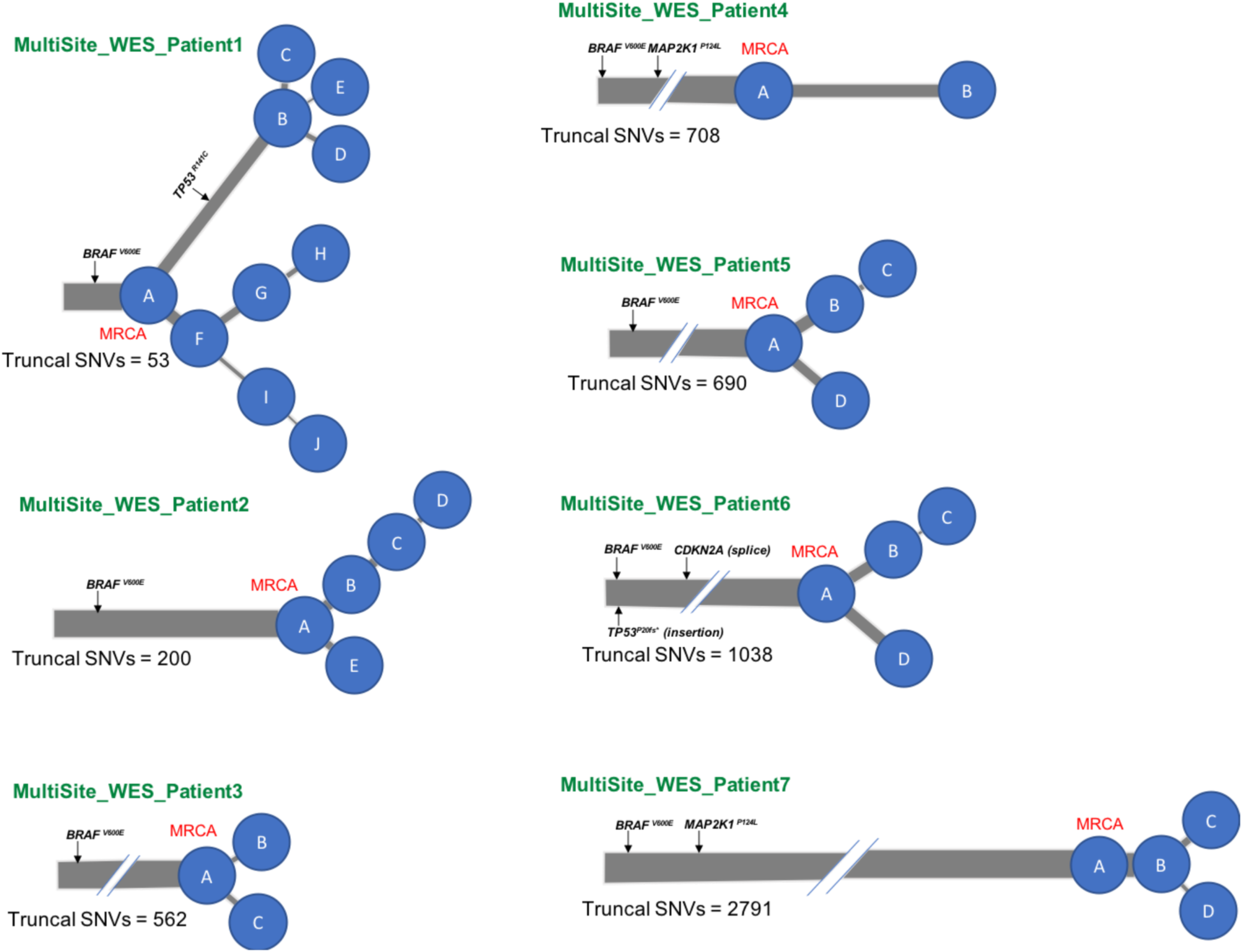
Multi-dimensional Dirichlet processing across metastases from a further 7 patients uncovers pervasive branched evolution. Phylogenetic trees of 7 metastatic melanoma patients are indicated, with branch lengths proportional to the number of SNVs within the mutation cluster and branch thickness proportional to the average CCF of the cluster (across all the samples within each patient). The number of truncal SNVs is indicated underneath the respective phylogenetic trees. All patients harboured an activating *BRAF^V600E^* driver mutation. In keeping with previous reports^13,15^, all other melanoma drivers were represented on the trunk (rather than the branches) of the phylogenetic trees, except for one patient harbouring a novel *TP53^R141C^* mutation which was unique to one branch and completely absent in the three samples on the opposing branch (despite adequate coverage, >40x).

### UV-induced mutational signatures were replaced by APOBEC-associated mutations across branches of the evolutionary tree

We extracted mutational signatures from the 13 whole-genome sequenced metastases collected from the index case as previously described^28^. As expected, all samples were dominated by signature 7 reflecting UV-induced mutagenesis (**Supplementary Fig. 5**). Within the pool of 2247 non-truncal mutations, however, we found evidence of non-UV induced mutational signatures, including signatures 2 and 13, which represent the action of the APOBEC family of cytidine deaminases (which enzymatically modify single-stranded DNA)^29^ (**Supplementary Fig. 6A**). We found that whilst signature 7 dominated the truncal cluster (and was also detected in the most recent common ancestor (MRCA) representing the earliest non-truncal mutational cluster), it was absent from the branches of the evolutionary tree, which were characterised by the APOBEC mutational signatures suggesting this process might be implicated in later stages of clonal evolution (**Supplementary Fig. 6B**). Interestingly skin cancer has been shown to have the fifth highest *APOBEC3B* expression rank^30^. However, the dipyrimidine-focused C-to-T mutation pattern of UV eclipses an *APOBEC3B* deamination signature, which we have only uncovered here by separating out the truncal mutations.

### Gene expression and immune cell estimation analyses show clustering of metastases within respective organs

Gene expression analyses of 11 metastases from the index autopsy case further revealed differences between metastases found in different organs, with principal component analysis (PCA) separating metastases sampled from the brain and lung. Gene expression profiles were also different between tumour and normal tissue from the same organ, indicating that these expression patterns most likely represent changes in tumour cell expression rather than organ-site related differences, suggesting an impact of the microenvironment on tumour gene expression^31^ (**Fig. 4A & Supplementary Fig. 7A & 7B**). Interestingly, metastases seeding within the brain clustered together by PCA, despite phylogenetic inferences indicating these likely emanated from differing lineages (**Supplementary Fig. 2**). By intersecting genes and pathways differentially expressed between brain metastases (n=5) and normal tissue (normal samples extracted from the brain and lung, n=2) (see **Methods** & **Supplementary Fig. 7C**), and also between the brain (n=5) and lung metastases (n=4) (**Supplementary Fig. 7D**), we identified particular genes and biological processes uniquely associated with brain metastases in this patient (**Fig. 4B**). The gene *PLEKHA5* was significantly up-regulated (log-fold change 4.5, FDR-adjusted p-value < *0.003*) in brain vs lung metastases, as well as in brain metastases vs normal (from the patients’ normal brain and lung tissue) (log-fold change 5.3, FDR-adjusted p-value < 0.004). This guanine nucleotide exchange factor has previously been shown to be upregulated in a cell line model of melanoma brain metastasis (cerebrotropic A375Br cells versus parental A375P cells) and silencing of *PLEKHA5* expression decreased *in-vitro* potential for crossing the blood-brain barrier^32^. Gene set enrichment analyses also showed significant enrichment of the KEGG pathway oxidative phosphorylation in both brain vs lung metastases (normalised enrichment score 4.65, FDR-adjusted p-value <0.0001) and in brain metastases vs normal (normalised enrichment score 3.17, FDR-adjusted p-value <0.0001) (**Fig. 4B**). This is consistent with recent analyses implicating the upregulation of oxidative phosphorylation in patient-matched brain versus extracranial metastases, as well as further functional studies demonstrating that inhibition of this pathway resulted in increased survival in both implantation xenografts and spontaneous murine models of melanoma brain metastases^33^.

**Fig. 4.**
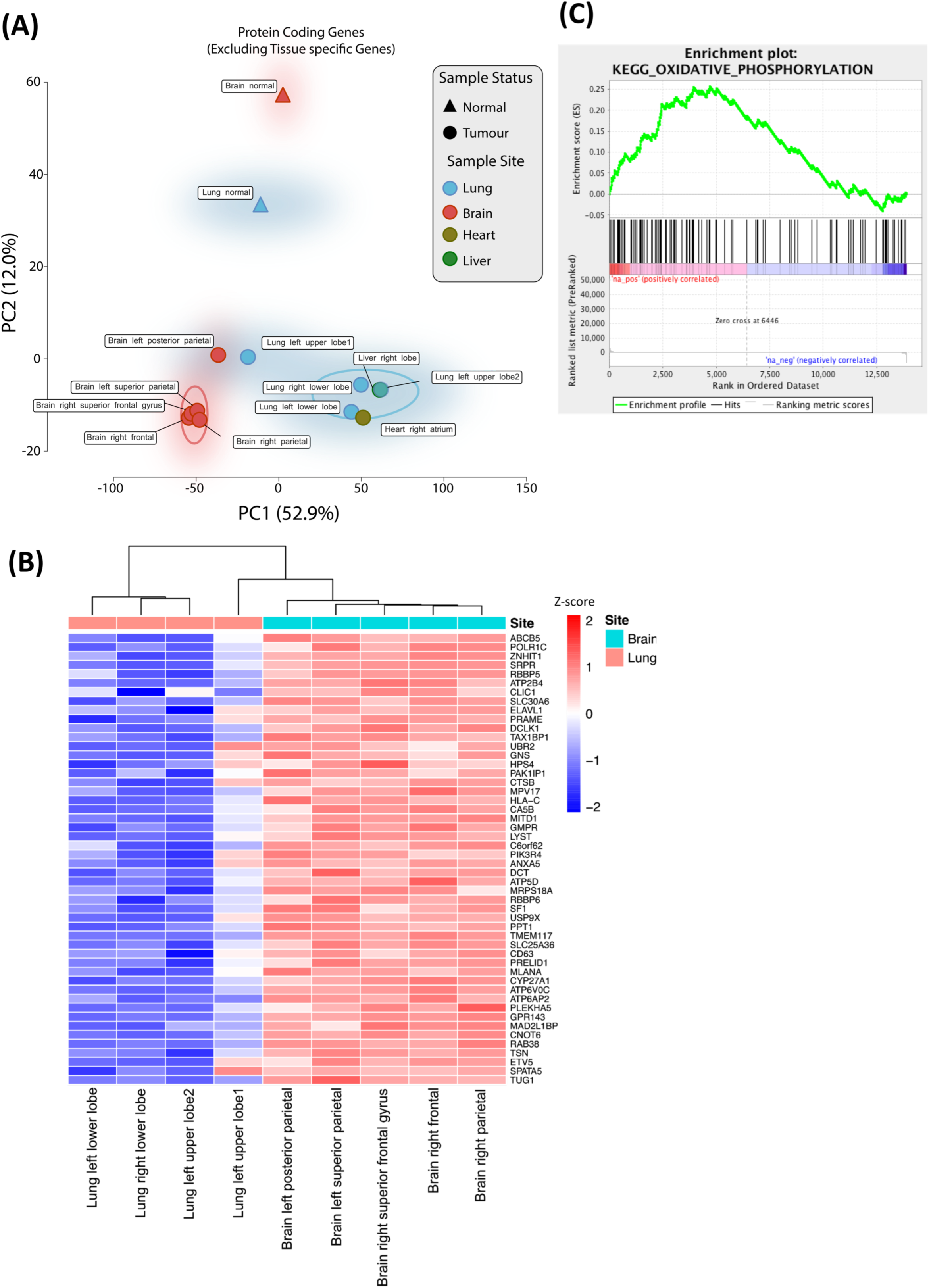
Gene expression analyses reveals regional separation of site-specific metastases, as well as genes and pathways that might predispose to brain metastases. A) Principal component analysis of protein-coding gene expression across all samples. A regional separation can be seen between the brain and lung metastases, which in-turn separated from the corresponding patient-matched normal organ control samples. The separation of tumour and normal samples indicates that the regional separation seen between brain and lung metastases is less likely to be confounded by non-tumoural cells. Samples are circled using a kernel density estimation. B) Heatmap showing the top 50 intersecting genes that are differentially expressed between both ‘brain vs lung metastases’ and ‘brain metastases vs normal’ comparisons (see **Methods**). Z-score indicates normalised gene expression and represents the number of standard deviations away from the mean. C) Preranked gene set enrichment analysis of genes associated with metastases to the brain revealed biological pathways that might have been implicated in brain metastases. The enrichment plot provides a graphical view of the enrichment score for a gene set. The top portion of the plot shows the running enrichment score for the gene set as the analysis walks down the ranked list. The middle portion of the plot shows where the members of the gene set appear in the ranked list of genes and the bottom portion of the plot shows the value of the ranking metric as you move down the list of ranked genes. Oxidative phosphorylation was the most statistically significant over expressed KEGG pathway in both the ‘brain versus normal’ and ‘brain versus lung’ comparisons, and has recently been linked to melanoma brain metastases in both human and murine analyses^33^.

Immune cell estimation of bulk tumoural mRNA from 11 of the metastases obtained from the index case (as previously described^34^), further revealed evidence of distinct tumour-immune microenvironments by immune cell representation (**Fig. 5**). The brain metastases in particular had relatively fewer immune cells compared to lung and other extracranial metastases, which might corroborate studies suggesting the brain represents a relatively ‘immuno-privileged’ organ^33,35^. Inflammatory macrometastases in the brain were however high for activated M2 macrophages (including significant up-regulation of the macrophage marker *CD163* in brain vs lung metastases, log-fold change 5.5, FDR-adjusted p-value < *0.004*) which have been described as having anti-inflammatory or tumour supporting activities, including in malignant brain tumours (although it is important to recognise that it may be difficult to distinguish between microglia and macrophages, both of monocyte lineage, using these methods)^36^. In summary, therefore, despite having overall similar mutational landscapes, we identified distinct tumour-immune microenvironments by gene expression, which likely reflects differences in the tumour microenvironment ecosystem^37^.

**Fig. 5.**
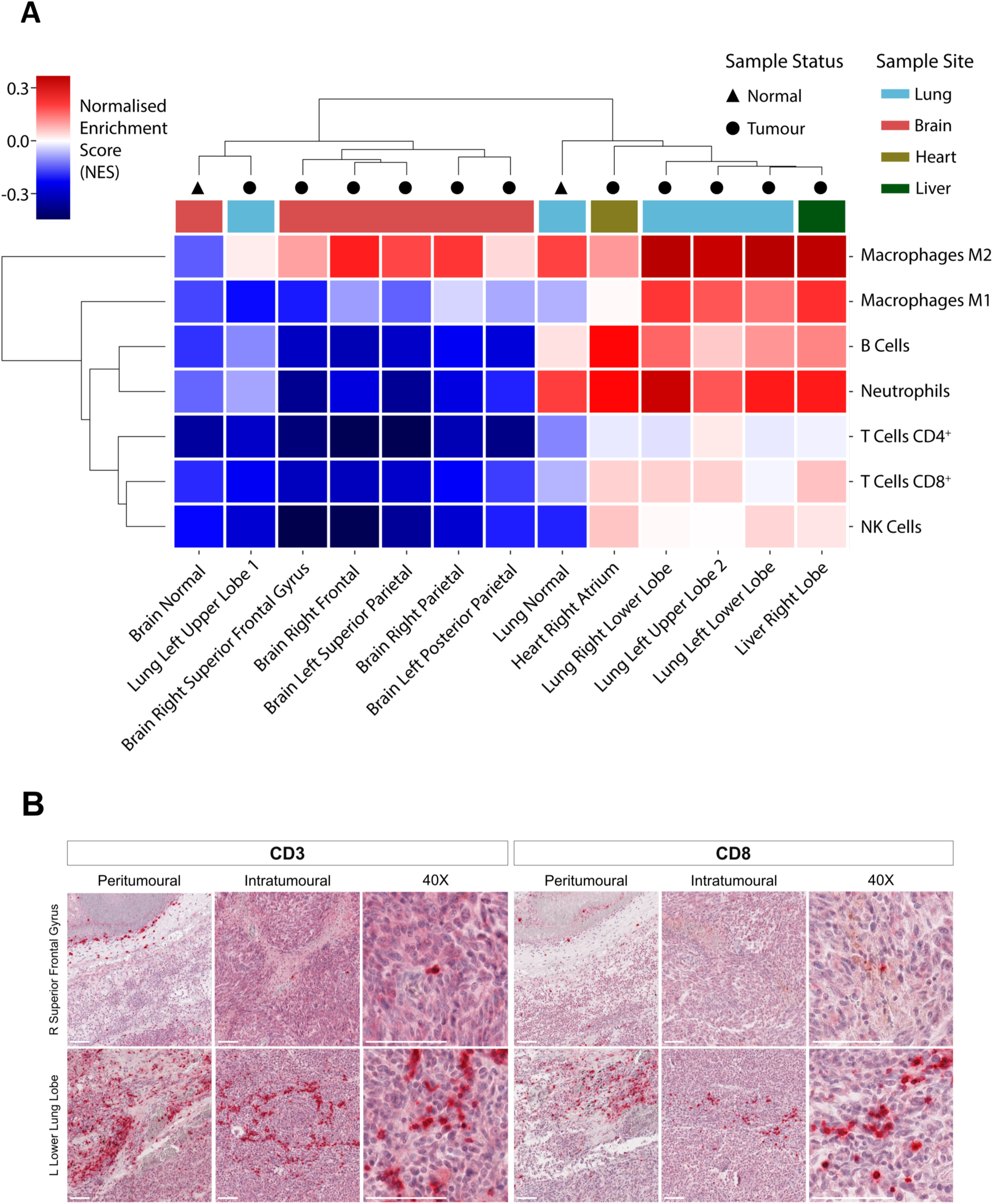
Regional differences in immune cell representation, with brain metastases being relatively ‘immuno-privileged’ relative to lung metastases. A) Immune cell deconvolution of mRNA using hierarchical clustering of Consensus^TME^ scores^34^. Both the brain normal and metastases were relatively ‘immuno-privileged’ when compared to lung normal and lung metastases (note that one of the four lung metastases, left upper lobe 1, was an outlier clustering with the immune-sheltered brain samples). B) Immunohistochemical (IHC) validation against CD3 and CD8 in a representative brain and lung metastasis. Chromogen red indicates immune cell staining. The thick white line represents 100um in 10-fold (peri- and intratumoural) and 40-fold magnification respectively.

## Discussion

Although melanoma is associated with a large number of somatic SNVs, the genomic diversity of melanoma metastases has previously been reported to be low, with most SNVs expected to be shared across tumours^13^. Recently, the International Cancer Genome Consortium Pan-Cancer Analysis of Whole Genomes (PCAWG) initiative^38^ leveraged WGS data to infer evolutionary relationships across multiple cancer types, further showing that metastatic melanomas may be monophyletic, with a single clone appearing to seed metastases and, when compared to other cancers, may uniquely lack subclonal heterogeneity^39^. In our study, longitudinal analyses of clone abundance from multi-site genome sequencing provided a powerful method to detect clonal selection and a unique insight into clonal evolution. In agreement with previous literature, we identified a single cluster of truncal variants ubiquitously represented across all metastases and representing >90% of all somatic SNVs. Truncal variants occurred early in tumour evolution and dominated downstream phylogenetic reconstruction analyses. Initial analyses therefore showed that subclonal heterogeneity appeared to be absent and that metastases were derived from a single parental clone harbouring the majority of genetic alterations. However, by subtracting this dominant cluster of variants, we were able to identify non-truncal clones and subclones. Assessment of the representation of these clones across the metastatic tumours revealed the ancestral relationships between metastases and, for the first time, uncovered conclusive evidence of branching evolution of metastases. In particular, these patient-matched primary and metastatic tumours provided the power to detect SNVs that were present only in a subset of sequenced samples, thereby increasing the power of these reconstruction approaches.

We found evidence of branched evolution across metastatic melanoma exomes from a further 6 out of 7 patients (including in one case of metastases from an acral primary), indicating that, even with a much lower sequencing depth and coverage, and nearly two orders of magnitude fewer SNVs (relative to whole-genome sequencing), detailed clonal lineages could still be inferred, and branching evolution is pervasive. The detection of branched evolution using lower-resolution WES from archival formalin-fixed paraffin embedded (FFPE)-derived samples is particularly relevant to clinical practice, where the majority of samples are still stored in paraffin, and where custom pull-down is much more readily available than whole-genome sequencing approaches^40/40^. It is therefore our impression that previous interpretations proposing linear evolution of melanoma metastases (where it has been thought that genetically distinct cell populations in the primary tumour might metastasise sequentially from one site to the next^13,23,39^), may have been confounded either by the use of VAF as a surrogate for CCF, or by the lack of power to separate subclones through single sample analyses (rather than by the limits of resolution of targeted sequencing approaches employed by many of these studies). In a previous single patient WGS study analysing a primary acral melanoma and its concurrent ipsilateral inguinal lymph node, a wide spectrum of SNVs and copy number alterations were found to be shared between the primary and metastatic tumour however, the phylogenetic architecture could not be fully reconstructed^41^. By harnessing the power of CCF calculations across 6 metastases from our acral melanoma patient, we identified branching evolution. A recent detailed multi-regional clonality analysis in uveal melanomas has also found multiple driver mutations in late branches of the phylogenetic trees, suggesting that these melanomas also continue to evolve as they progress from primary to metastatic disease^42^. Therefore, we postulate that branched evolution is characteristic of all melanoma subtypes.

Analysing skin/subcutaneous metastases in 8 patients with cutaneous melanoma, Sanborn *and* colleagues previously showed that locoregional relapses arose from different subpopulations of the primary tumour cells, which often disseminated in a parallel rather than serial fashion^43^. Our phylogenetic analyses support these findings, and show that branched evolution is associated with both locoregional as well as more distant metastatic spread. Our finding that truncal mutations were identified as a subclone within the index patients’ cutaneous primary further corroborate the observation that metastases likely seeded early, at a time when distant disease was clinically undetectable. Interestingly, the brain metastases from the index autopsy case were represented across both branched lineages (right parietal and right frontal brain metastases harbouring subclonal mutations represented across distinct clonal lineages **Supplementary Fig. 2**), suggesting there were at least two waves of spread (polyclonal seeding) to the brain. However, in keeping with previous analyses^15,43,44^, polyclonal seeding was generally a rare event in this cohort (**Supplementary Fig. 2 & 4**). Although the index patient underwent whole-brain radiotherapy six weeks prior to the autopsy, we did not detect differences in mutational signatures between the brain and extracranial metastases^28,45^ (**Supplementary Data**), while the lack of prior systemic therapy further supports the mutational processes being reflective of evolutionary changes during dissemination, rather than drug-induced.

The phylogenetic trees were characterised by non-truncal SNVs appearing late in the evolutionary course and represented by rapid branching of the phylogenetic tree from a long trunk (‘palm tree’ resemblance) (**Fig. 2D**). Similarly, driver mutations generally arose before subclonal diversification and were found primarily on the lung trunks of the trees (**Fig. 2D & 3**). This contrasts with what has traditionally been thought to be a slow iterative process of gradual evolution, as typified by recent reports in prostate cancer^9^, where branching generally occurred earlier and more gradually throughout the tumours’ evolutionary trajectories, as well as driver mutations being frequently observed subclonally (rather than predominantly clonally in this and other analyses^13,15^). Although the index patient was initially diagnosed with a low-risk stage IB cutaneous primary, predicted to have >95% 5-year survival^46^, the time from detection of metastatic spread to death from disease was very short, which is not uncommon and contributes to the challenges of managing this disease. Further prospective studies will be required to confirm our findings, suggesting a short latency between emergence of the invasive clone and widespread metastases, and to determine how they interplay with the established melanoma prognostic markers^46^.

The catalogue of somatic mutations in cancer is the aggregate outcome of exposure to one or more mutational processes. Each process generates mutations characterised by a specific combination of nucleotide changes and nucleotide contexts, therefore providing a signature that can be used for its identification^28,47^. Mutations in cutaneous melanomas are dominated by C>T transitions caused by mis-repair of ultraviolet (UV) induced covalent bonds between adjacent pyrimidines (signature 7)^1^. Studies by Shain and colleagues suggested that UV is the dominant factor associated with the initiation of precursor lesions and dominates every stage of tumour evolution, from the progression of pre-malignant lesions to primary melanoma and to metastases^16,17^. Interestingly however, other studies have reported a reduction of the proportion of mutations associated with the UV-induced mutational signature in branch (non-truncal) mutations, suggesting that mutations arising later in melanoma progression may occur as a result of increased activity of other mutational processes^13,15^. This is consistent with lung cancer analyses, where subclonal lineages acquired mutations that lacked the tobacco-smoking signature and were replaced with mutations associated with APOBEC cytidine deaminase activity^7^. In keeping with these analyses, we found that whilst the truncal cluster of mutations in the autopsy index case was dominated by signature 7, this was completely lost in the subclonal mutation clusters which also appeared to be replaced by APOBEC-associated mutations. Further studies will be required to ascertain whether these can unravel new biological insights.

In summary, through leveraging the power of clonality analyses across multiple whole-genomes we were able to identify rich clonal architectures and uncover pervasive branched evolution of melanoma metastases obtained at autopsy of a single patient, a structure which would not have been evident through single-site reconstructions. This pattern of phylogenetic branching was also evident in exome sequenced metastases obtained from 6 out of 7 additional melanoma patients, one of which was an acral melanoma, suggesting that this is independent of sequencing coverage, depth, the number of SNVs, or melanoma subtype. Our ability to detect distinct clonal lineages was greatly enhanced by leveraging the power from multiple samples. Our data reframes current models of metastatic dissemination and should serve as a cautionary tale in future phylogenetic analyses that define trunk and branch mutations by the presence or absence of shared variants and that do not consider CCF calculations (integrating information from somatic VAFs with tumour purity and ploidy considerations). Future large-scale studies incorporating clonal analyses across multiple metastases will be required to further delineate how these tumours evolve, and provide insights into whether interrupting this process could contribute to patient management.

---

**Supplementary Fig 1.**
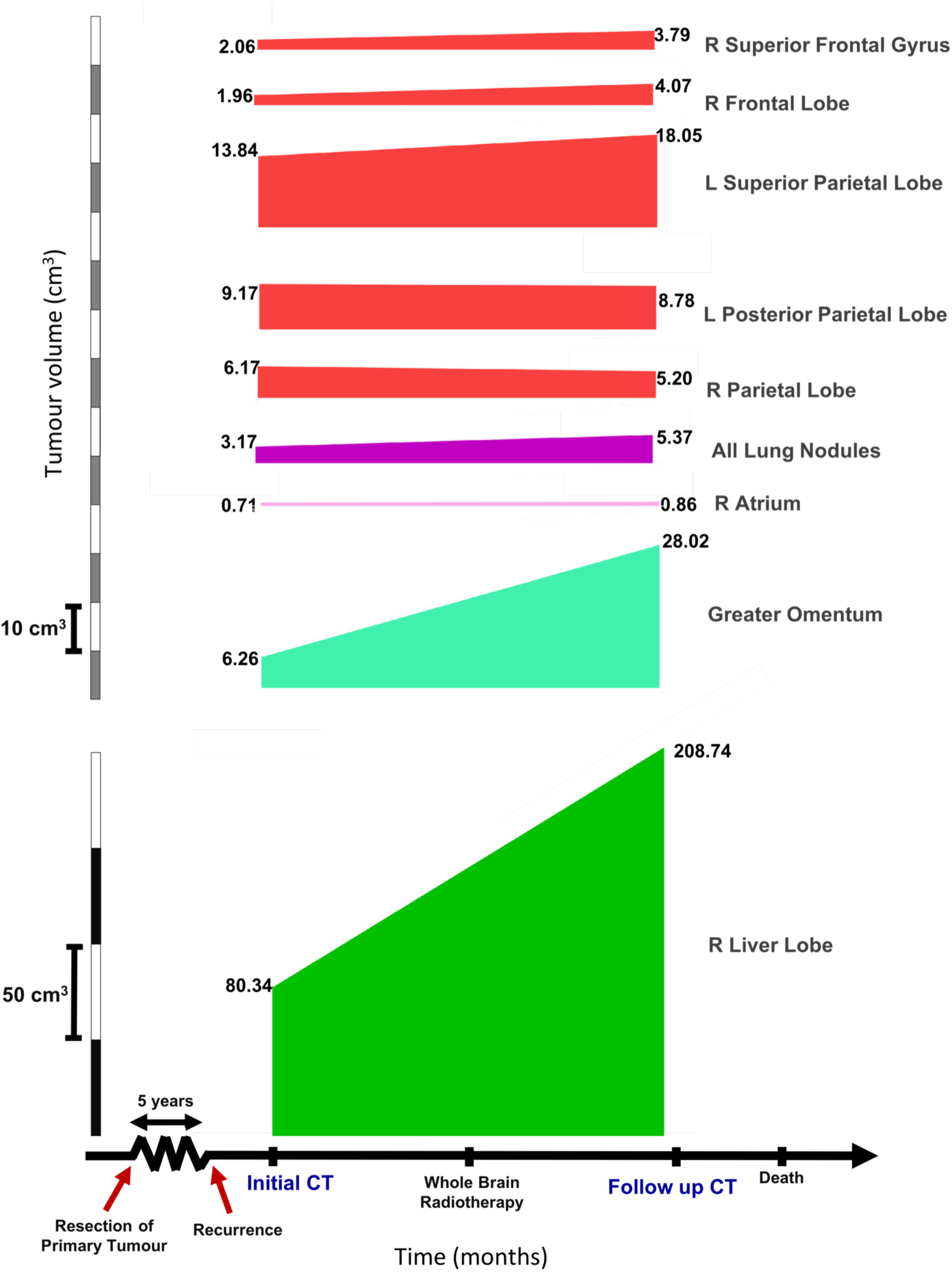
Representative clinical timeline of the index autopsy case, demonstrating rapid progression from the first appearance of metastatic disease. The volume changes of target lesions between interval CTs performed 5 weeks apart are shown, with the second imaging taken two weeks after the completion of whole-brain radiotherapy. This showed only minimal intracranial, but extensive extracranial disease progression.

**Supplementary Fig 2.**
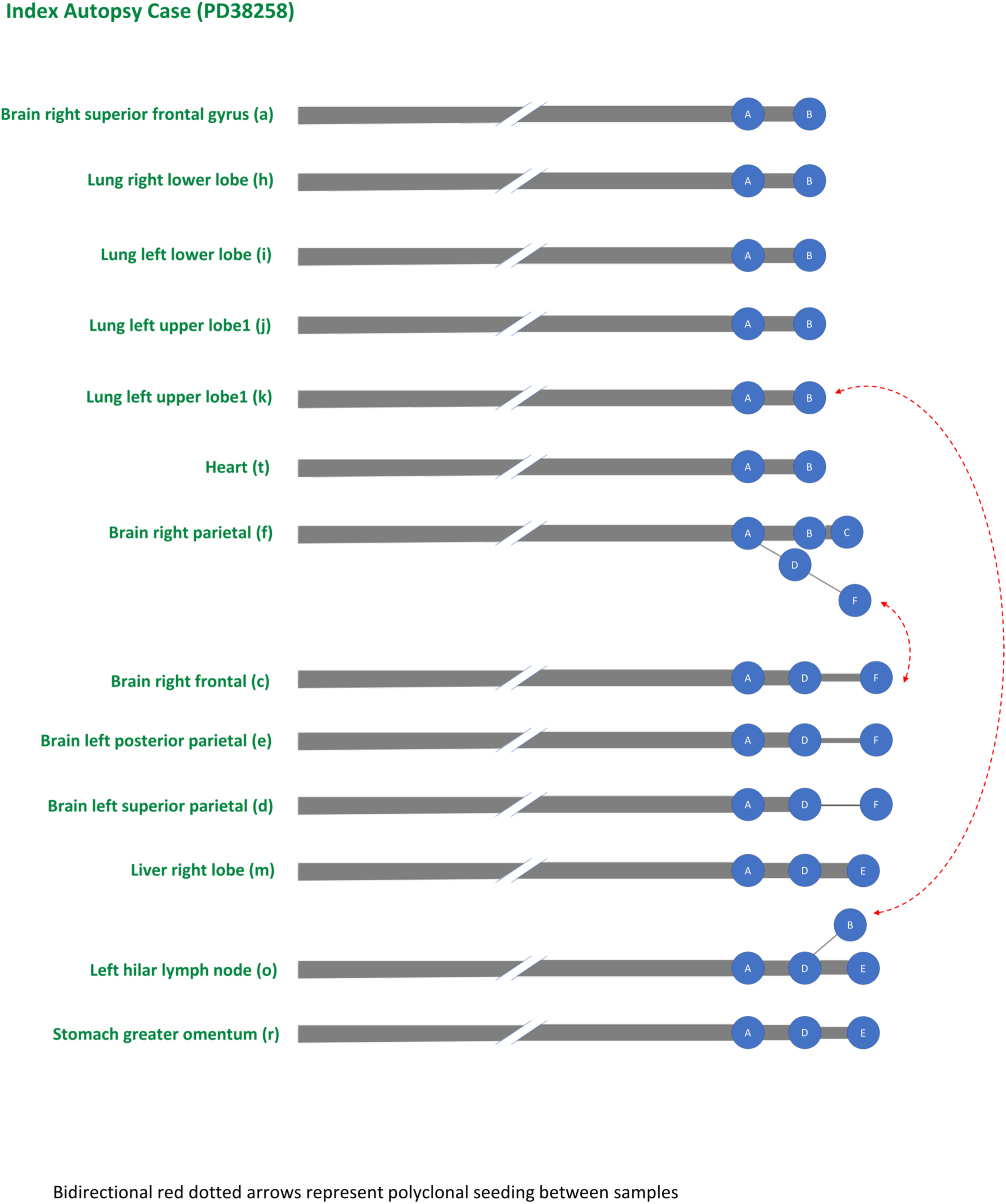
Sample-level phylogenetic tree for the index autopsy case. Each tree represents a subtree of the overall phylogenetic tree (Fig. 2D) including just those subclones seen within that particular sample, however in doing this we were able to segregate the samples based on their respective clonal lineage. We observed two clear lineages, representing distinct waves of metastatic seeding. Bidirectional red dotted arrows represent polyclonal seeding between samples. Interestingly, 4 metastases (2 from the brain, 1 from the lung and hilar lymph node respectively) from the index case displayed evidence of polyclonal seeding, being represented across both branched lineages.

**Supplementary Fig 3.**
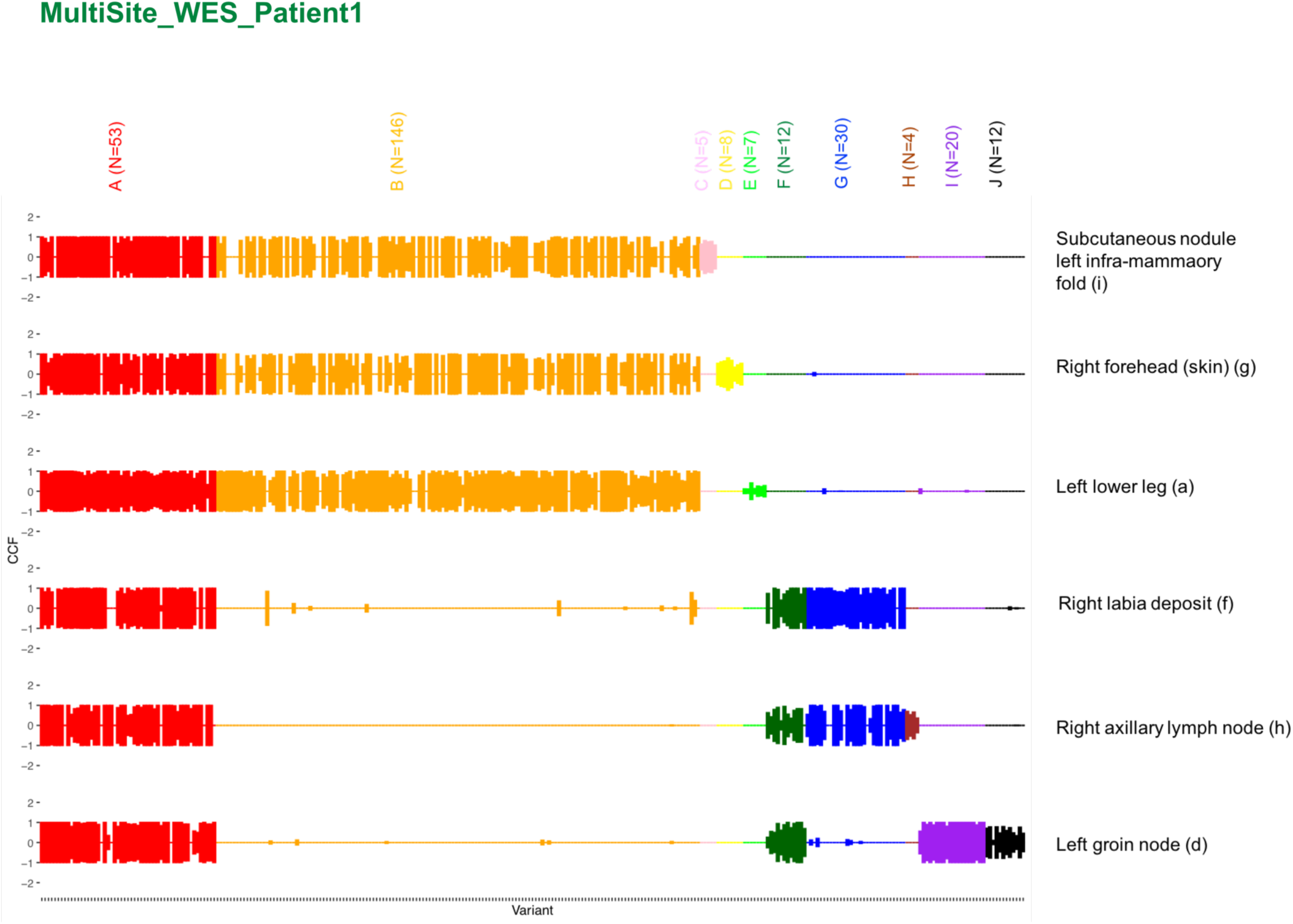
CCF distribution plot for whole-exome sequenced patient MultiSite_WES_Patient1. Rows reflect samples and columns reflect alphabetically and colour-coded mutation clusters (number of SNVs within each cluster is indicated at the top).

**Supplementary Fig 4.**
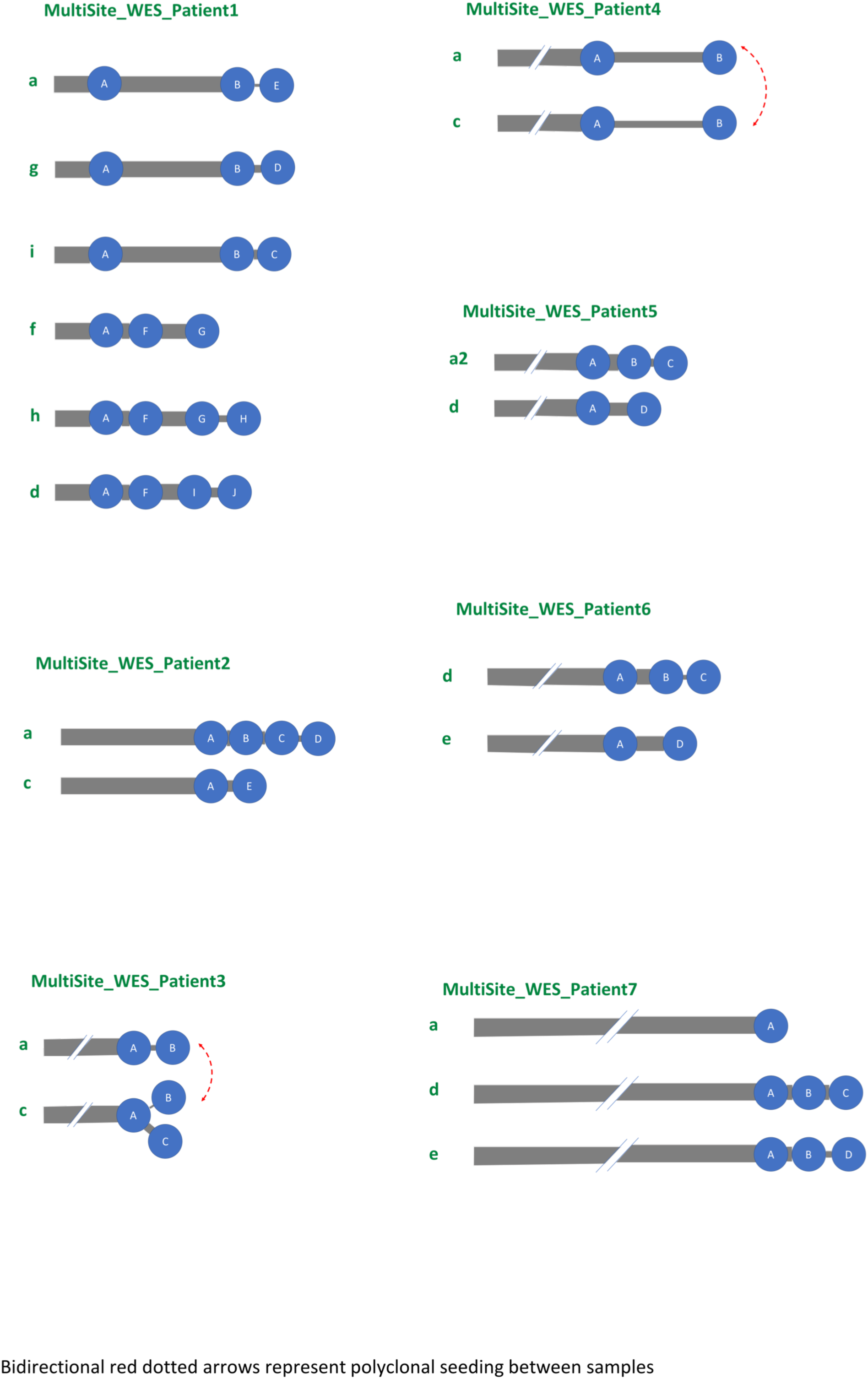
Sample-level phylogenetic tree for multi-site whole-exome sequenced cases. Bidirectional red dotted arrows represent polyclonal seeding between samples, which was observed in two cases (MultiSite_WES_Patient3 and MultiSite_WES_Patient4).

**Supplementary Fig 5.**
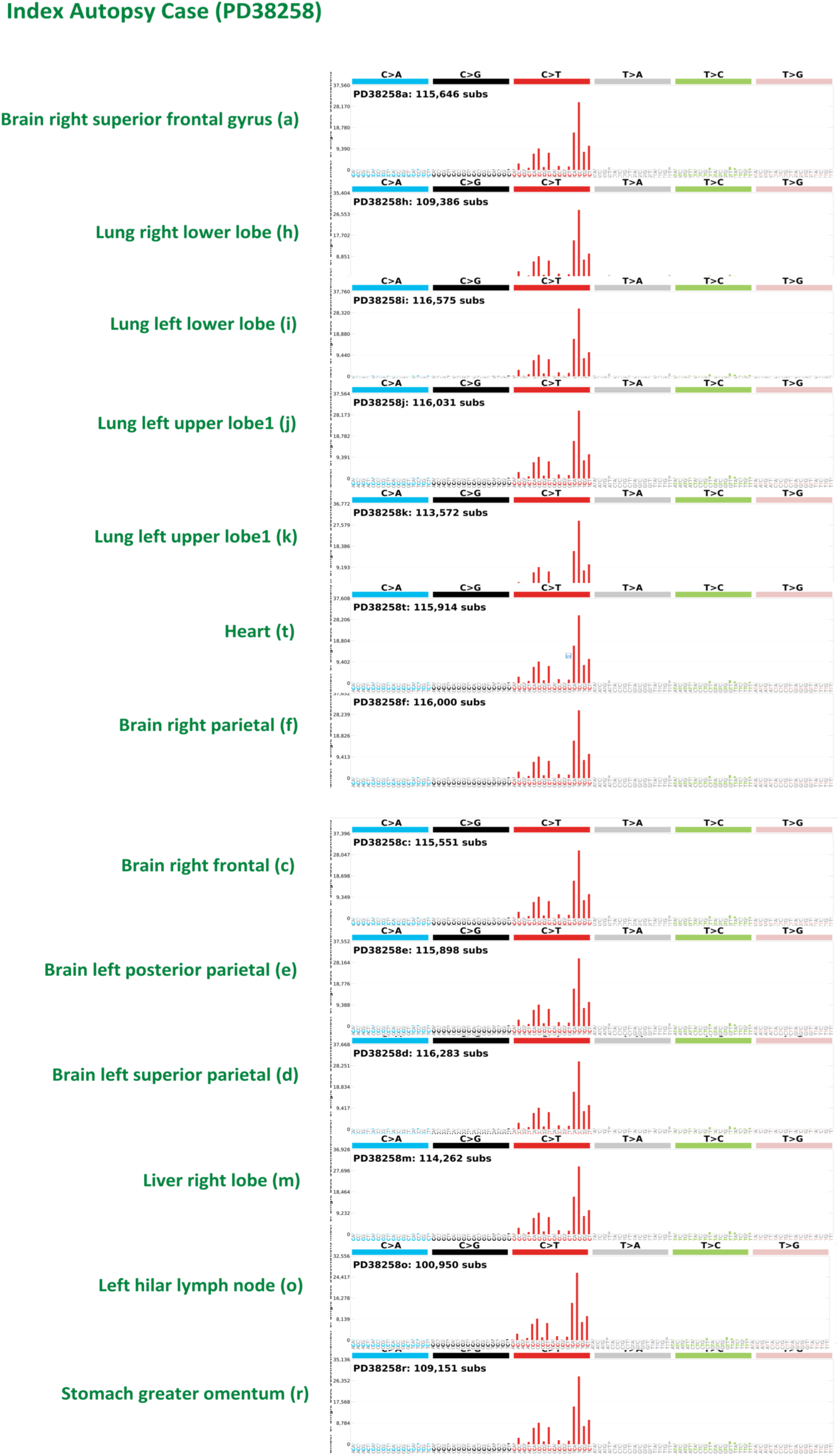
Mutational signatures for all SNVs from the index autopsy case. Shows the mutational profile using the conventional 96 mutation type classification as described by Alexandrov and colleagues^28,47^. This classification is based on the six substitution subtypes: C>A, C>G, C>T, T>A, T>C, and T>G. Further, each of the substitutions is examined by incorporating information on the bases immediately 5’ and 3’ to each mutated base generating 96 possible mutation types. Here we show the signature profiles including all SNVs from all 13 WGS metastases which as expected, was dominated by signature 7.

**Supplementary Fig 6.**
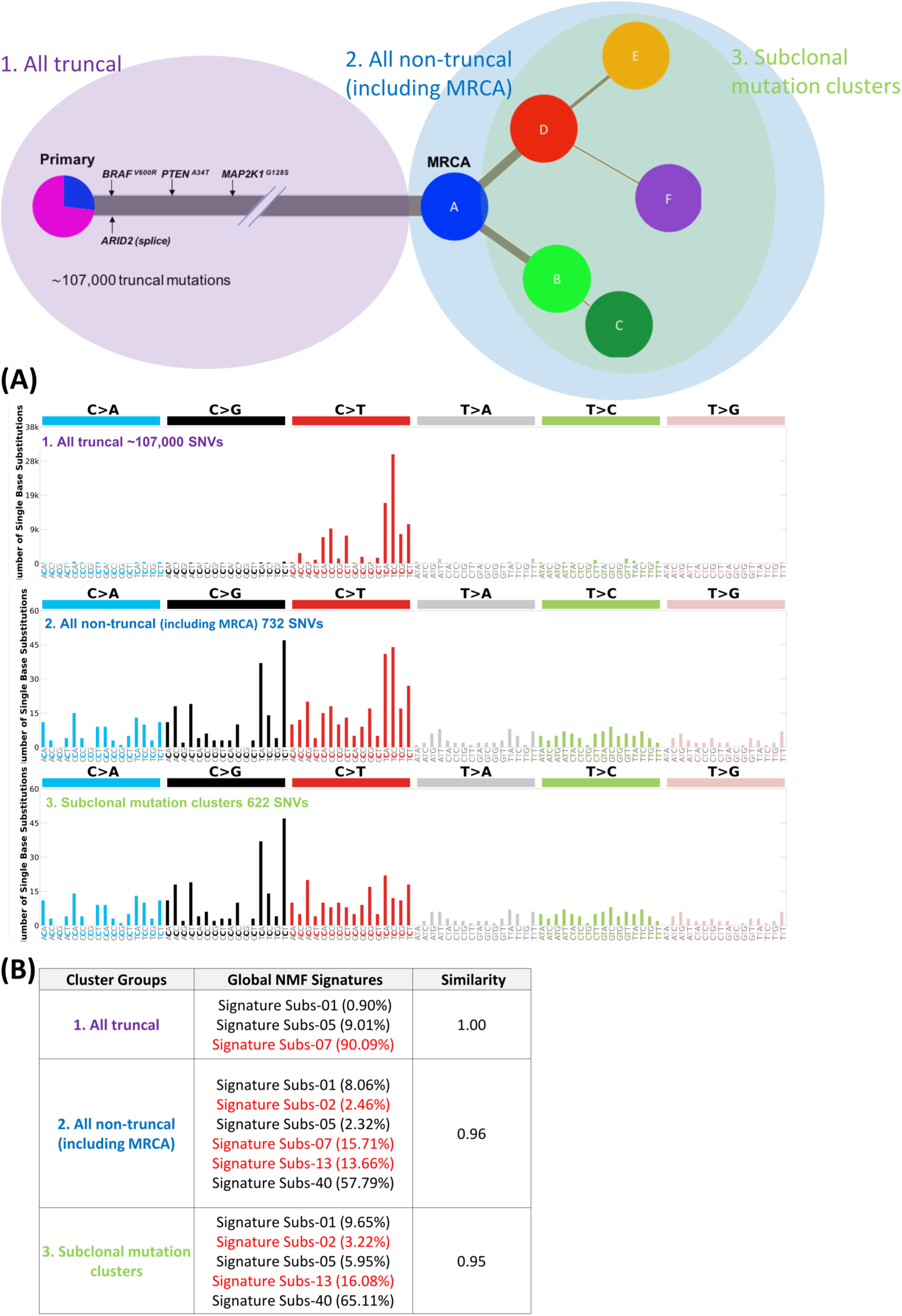
Mutational signatures from the index autopsy case. A) Here we show the mutational profiles including; all truncal SNVs in the tree (n∼107,000 SNVs), all non-truncal SNVs in the tree (including the MRCA, n=732 SNVs) and all subclonal mutation clusters in the tree (excluding the MRCA, n=622 SNVs). B) The Global NMF signatures shown represent the ‘best fit’ signatures across all SNVs and the individual percentages for each signature is the proportion which that signature represents. The cosine similarity reports how closely these signatures together mirror the context of all SNVs within that cluster group. As expected, the mutational signature for all truncal SNVs is dominated by signature 7 (90% of SNVs are represented by this signature), the non-truncal SNVs (including the MRCA) includes 15% representation from signature 7 whilst this is entirely absent from the subclonal branched lineages, which are represented by the APOBEC mutational signatures (2 & 13). The non-highlighted signatures (1, 5 and 40) shown in black represent relatively featureless (“flat”) oncogenic signatures found in most cancer types and do-not as yet define any distinguishing biological processes^47^.

**Supplementary Fig 7.**
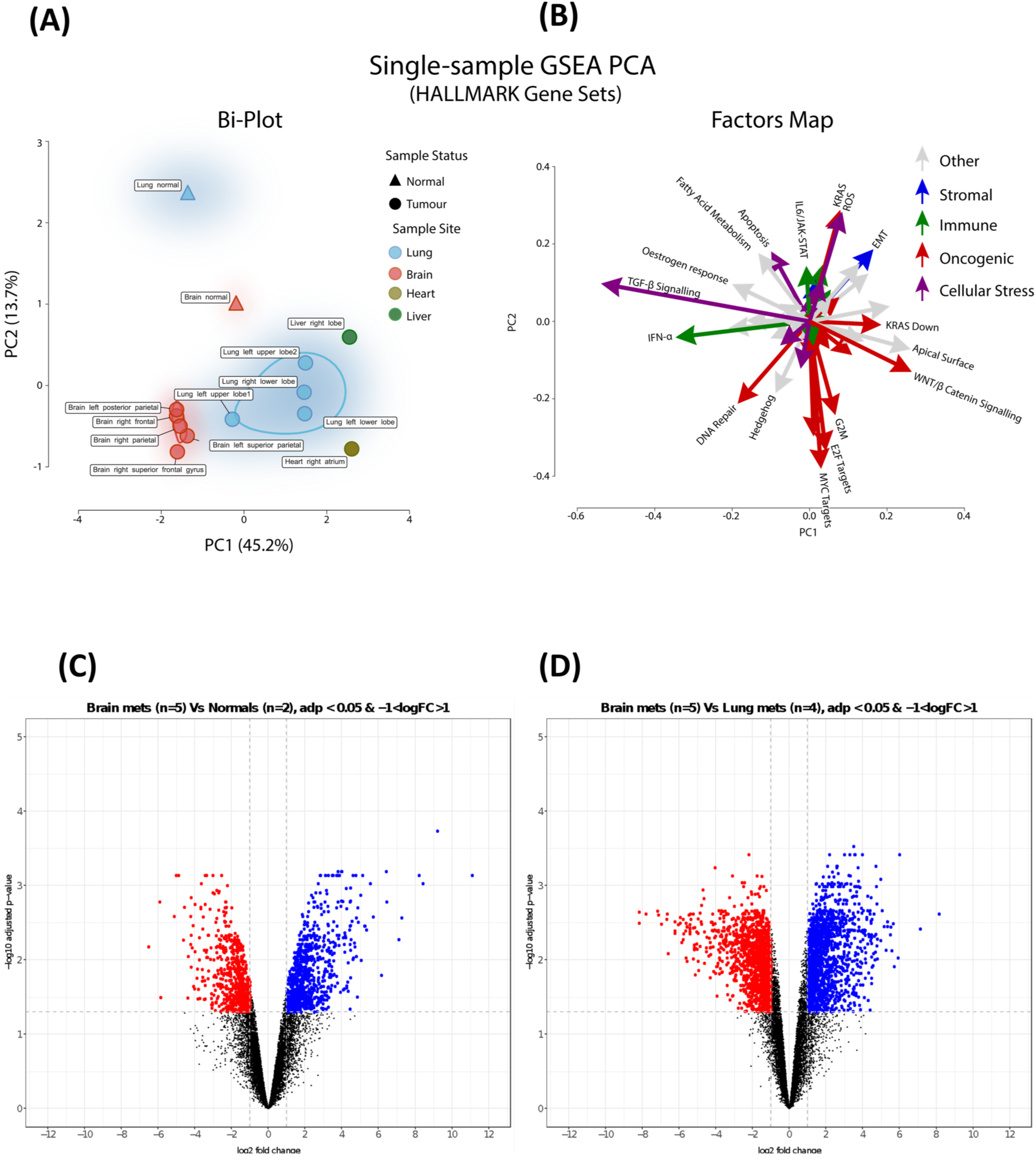
Differential expression and principal component analyses from the index autopsy case. A) Principal component analysis (PCA) of single sample gene-set enrichment (ssGSEA) using the hallmark gene sets^48^. A similar pattern of regional separation (as represented in Fig. 3A) was observed between the brain and lung metastases, which again are separated from the corresponding patient-matched normal organ control samples. Samples are circled using a kernel density estimation. B) Principal component feature loadings (magnitude and direction) of (A) are shown in the variables factor map. Each co-ordinate in (B) reflects the correlation coefficient of the biological process to principal components 1 (x-axis) and 2 (y-axis) from (A). Vectors are coloured according to the major biological classification of Hallmark gene sets. This revealed that PC2 (on the y-axis), explaining the variation between the tumour and normal samples (represented by circles and triangles in (A) respectively), is primarily represented by the up-regulation of oncogenic processes highlighted with a red arrow pointing (downwards) towards the tumour samples. C) Volcano plot of the genes differentially expressed between the brain metastases (n=5) versus the patient-matched normal tissue (one sample from brain and lung respectively, n=2). Each dot represents one gene and colours represent the significance (FDR-adjusted p-value <0.005) and fold-change cut-offs (log fold-change < −1 are coloured in red, and log fold-change >1 in blue). D) Volcano plot of the genes differentially expressed between the brain (n=5) versus lung metastases (n=4). Dots represents genes, coloured in the same format as (C).

**Supplementary Table.** Summary of key clinical details relating to patients within this study, including the sample IDs mapping to the raw sequencing files and variant/CNV calls, which are all deposited in the **Supplementary Data**.

## Methods

### Patient enrolment

All patients were recruited to the MelResisist prospective non-interventional study sponsored by Cambridge University Hospitals NHS Foundation Trust. The study was approved by the National Research Ethics Committee (NREC) North East on the 17^th^ October 2011, IRAS project ID 66161 and REC reference 11/NE/0312. All patients provided written informed consent to take part. All cases were also ethically approved by the Sanger Institute’s human materials and data management committee. The research autopsy was conducted 48h after the patient’s death, during which time the body had been stored at 4°C. Sixteen 1cm-diameter core biopsies were sampled from the centre of each metastatic tumour and snap frozen in liquid nitrogen at −80°C. These were used for the extraction of bulk DNA and RNA, as well as for the creation of stained H&E slides for direct histopathological assessment of the sequenced regions. The remaining multi-site exome-sequenced cases were also identified from the prospective MelResist trial, and were selected based on their availability of banked multi-site metastases for molecular interrogation. A total of 7 patients with 21 metastases (median 2, range 2-6 metastases per patient) were identified for this analysis. All samples and clinical details are listed in the **Supplementary Table**.

### Extraction and quality assessment of DNA and RNA

Histopathological assessments were performed by two consultant histopathologists (KA and MT), who confirmed that all tumours were composed of >90% neoplastic cells. Macro-dissection of fresh tumour cores from the autopsy case was performed with a sterile scalpel. DNA and RNA were extracted from the fresh tumour cores using the AllPrep combined DNA/RNA Mini Kit (Qiagen Ltd.) according to manufacturer’s recommendations. All the multi-site exome-sequenced cases were obtained as 1.0mm diameter cores micro-dissected from the original FFPE block. Genomic DNA was extracted from the FFPE cores using the QIAamp FFPE Tissue kit from Qiagen according to manufacturer’s instructions. Germline DNA was extracted from peripheral blood mononuclear cells collected before death from all cases, using the DNeasy Blood and Tissue Kit (Qiagen). To confirm that the tumours and germline DNA were derived from the same patient, genotyping was performed using 96 SNP6 (Fluidigm) and confirmed that all the samples were derived from the same individual. All DNA samples were quantified using the PicoGreen dsDNA Quantification Reagent according to manufacturer’s recommendations (Invitrogen). The structural integrity of DNA was checked by gel electrophoresis. RNA quantity and quality were assessed using Agilent’s 2100 bioanalyzer.

### Laser capture microdissection of the cutaneous primary from the autopsy case

Two FFPE tissue blocks from the index autopsy patient’s archival primary tumour (cutaneous melanoma from the anterior chest wall, samples PD38258u and PD38258v) were processed into 5µm histology sections, deparaffinised with ethanol thrice and stained with Gill’s haematoxylin for 20 seconds. Malignant melanocytes from each section were isolated by a histopathologist (LM) using laser capture microdissection and collected in separate Eppendorf tubes. These were then lysed with lysis buffer ATL and digested with proteinase k (Qiagen Ltd.). Extraction of nucleic acids was performed using the QIamp DNA FFPE extraction kit (Qiagen Ltd.) according to manufacturer’s recommendation.

### Whole genome sequencing and somatic variant detection

Paired-end sequencing of the metastatic tumours and matched normal was performed on the Illumina X10 platform at the Wellcome Trust Sanger Institute to generate 150 base-pair reads. Sequencing reads were aligned using BWA-MEM (v0.7.12)^49^ to the human reference genome (NCBI build GRCh37). The resulting sequencing coverage ranged from 33- to 43- fold (median 38- fold). Caveman (v1.11.2)^50^ and Pindel (v2.2.4)^51^ were used to call somatic SNVs and indels, respectively. The minimum base quality score for somatic variant calling was set to Phred 30. ANNOVAR^52^ was used to predict the effect of variants on genes and to assign rsIDs for known variants based on dbSNP Human Build 150. The alignments for all variants are reported in **Supplementary Data**. To call rearrangements we applied the BRASS (breakpoint via assembly) algorithm, which identifies rearrangements by grouping discordant read pairs that point to the same breakpoint event (github.com/cancerit/BRASS). BRASS rearrangements were used to search for balanced inversions, which have been previously associated with radiation-induced mutagenesis^45^, and were particularly relevant to explore in the radiotherapy-treated brain metastases from the index case.

### Whole exome-sequencing of multi-site metastases cases

Exome capture was performed using Agilent’s SureSelect bait. Paired-end sequencing was performed using the Illumia HiSeq platform at the Wellcome Trust Sanger Institute to generate 75bp reads. Sequencing reads were aligned using BWA-MEM (v0.7.12)^49^ to the human reference genome GRCh37. PCR duplicates, secondary read alignments, and reads that failed Illumina chastity (purity) filtering were flagged and removed prior to running variant and copy number calling. The resulting sequencing coverage after filtering ranged from 33- to 95-fold (median 52-fold) in the tumoural samples and median 54-fold across the germline blood samples. Caveman (v1.11.2) and Strelka (v1.0.15)^53^ were used to call somatic SNVs and indels, respectively. The minimum base quality score for somatic and germline variant calling was set to Phred 30. ANNOVAR^52^ was used to annotate SNVs (based on Caveman) and indels (based on PINDEL) for functional classification and to assign rsIDs for known variants based on dbSNP Human Build 150 (**Supplementary Data)**.

### Copy number aberration (CNA) profiling

Segmental copy number information was derived for each of the 13 metastatic tumours using the Battenberg algorithm (v3.2.2)^8^. This was also used to estimate tumour cellularity and ploidy, and calculate allele-specific copy number profiles as previously described^10^. Sequenza (v2.1.2)^54^ was used to estimate tumour cellularity and ploidy from the tumour-normal pairs in the multi-site FFPE-extracted exome sequenced cohort, as well as to calculate allele-specific copy number profiles (**Supplementary Data)**. For each sample, the best Sequenza solution was chosen after visual inspection of both the best-fit solution (with the maximum log posterior probability) and alternative solutions.

### Validation of SNVs from the whole-genome sequencing analysis

Validation was performed using custom pull-down and sequencing of the mutations identified in the WGS analysis. The validation experiment was enriched to cover all 2247 non-truncal variants, 652 manually selected truncal variants (identified as either cancer driver mutations or with loss-of-function mutations from the truncal cluster) as well as 10 shared frameshift variants common to all metastases. A 340kbp custom capture probe was designed using Agilent Technologies’ online software ‘Sure Select Design Wizard’. The highest-stringency repeat masking was used (where possible), as well as a tiling density of 2X and maximum performance boosting (replicating any orphan or GC-rich baits by a higher factor). DNA capture (paired-end, average DNA fragment size 158bp) libraries were created using native DNA, testing DNA from all 13 whole-genome sequenced metastatic tumours. Libraries were multiplex sequenced to a median depth of 40X on the Illumina MiSeq platform. A variant called in the WGS experiment that was also present in the validation study and supported by at least 2 alternate bases in the validation, is reported as validated somatic. With these criteria, 7429/7502 (99%) of the truncal substitutions and 6223/6750 (92%) of the non-truncal substitutions were validated somatic (the denominator represents the sum of all the SNVs called at each of the 13 samples in the WGS data, excluding those SNVs where coverage in the validation experiment was <30x, **Supplementary Data**). Only 7/7502 of the truncal mutation calls made in the WGS were not detected in the validation experiment, and 104/6750 (1.5%) of the non-truncal mutation calls were not detected in the validation experiment.

### Targeted sequencing of the archival primary

The same baits and custom pull-down experiment described above were used to sequence the validation set SNVs within the index autopsy case’s primary tumour. DNA was extracted from the two tumour blocks from the same chest wall primary and variants supported by at least 2 alternate bases were called in the primary (samples PD38258u and PD38258v).

### Gene expression analyses

RNA expression of metastatic tumours from the index autopsy case was determined using the human Affymetrix Clariom D Pico assay (**Supplementary Data)**. Arrays were analysed using the SST-RMA algorithm on the Affymetrix Expression Console Software. Expression was determined using the Affymetrix Transcriptome Analysis Console. Median absolute deviation normalization (of probe-level data) was implemented. The Tissue-specific Gene Expression and Regulation (TiGER) database was used to filter out 600 ‘tissue-specific’ genes^55^. Differential expression was performed using the R package limma (v3.36.1)^56^. Preranked GSEA (GSEA-P) was implemented using the GenePattern module *GSEAPreranked* (v6.0.10)^57^. Hallmark gene sets were downloaded from the MSigDB database^48^. Rank metric was calculated as the sign of log2-FCs calculated using the limma pipeline. Single sample gene set enrichment analysis (ssGSEA) was employed using the ‘GSVA’ R package (v1.32.0) to determine the relative enrichment of each of the HALLMARK pathways across samples^58^.

### Immune cell deconvolution

A novel consensus approach, Consensus^TME^ was used to generate cell-type specific estimates of immune cell infiltration from bulk tumour RNA gene expression profiles. This leverages information from multiple gene sets and immune cell expression matrices to build a compendium of robust gene sets for each immune cell type^34^. These genes were further filtered to ensure each has a negative correlation with tumour purity within The Cancer Genome Atlas. Gene sets specific for human skin cutaneous melanoma (SKCM) were used. Finally, single sample gene set enrichment analysis was applied to our Consensus^TME^ gene sets to generate normalised enrichment scores for each cell type in each sample. This method has previously been thoroughly benchmarked in a pan-cancer setting^34^.

### Extraction of mutational signatures

SigProfilerMatrixGenerator python packages^59^ was used to extract mutational signatures, generating 96 possible mutation types and used to plot mutational profiles. The sigproSS python package (v0.0.0.26)^60^ was used to determine the proportion of mutations in each sample attributable to specific COSMIC signatures identified by Alexandrov et al^28^. sigproSS was also run on the non-truncal mutation clusters defined in the phylogenetic tree (**Supplementary Fig. 5**).

### Computerised Tomography Analysis of Tumour Volume

Regions of interest were outlined over the entire area of visible tumours on post-contrast CT scans (2mm thin sections) by a radiologist (FS) using the OsiriX medical imaging software (Pixmeo SARL, Switzerland). Tumour volume was calculated by multiplying the area of tumour outlined on each CT image by the slice thickness.

### Immunohistochemistry

IHC staining was performed on 5µm FFPE sections, extracted from the same frozen sections from which DNA and RNA were extracted. Slides were deparaffinised in series of xylene and hydrated in a series of descending ethanol. Heat-induced antigen retrieval was performed using TRIS-EDTA (pH = 9), followed by immunostaining performed on the Leica Bond III autostainer (Leica Biosystems). Antibodies used included mouse monoclonal anti-human CD3 (DAKO, clone F7.2.38, dilution 1:50) and mouse monoclonal anti-human CD8 (DAKO, clone C8/144B, dilution 1:25) at 1 hour RT. DAKO REAL^TME^ alkaline phosphatase and chromogen red detection system was used for secondary detection of positive staining. Stained slides were counter-stained with haematoxylin and cover-slipped for review. Image acquisition was performed on the Hamamatsu whole slide scanner at 40-fold magnification.

### Statistical analysis and informatics approaches

All statistical analysis and graphics were generated using R version 3.0.1 (R Foundation for Statistical Computing, Vienna, Austria. URL http://www.R-project.org/). Alignment viewing was performed using Jbrowse and IGV.

### Analysis of Intra-tumoural heterogeneity (ITH) and phylogenetic tree reconstruction

To model the clonal structure across all multi-site tumour samples per patient (at WGS, WES and targeted sequencing levels), we used a previously described computational framework^61^. This approach is an SNV-centric ITH analysis which is briefly described below. In the first step, CCF is estimated for each SNV. By taking into account VAF, CNA status of the SNV locus and purity of the tumour sample under analysis, mutation copy number^62^, which is the product of CCF and number of SNV-bearing mutations, is calculated. CCF is then estimated from mutation copy number by adjusting for the number of SNV-bearing chromosomes, as assessed by a binomial distribution maximum likelihood test^61^. SNVs were removed from further analysis if they occurred in regions with different CNA status across samples and loss of heterozygosity or any other altered CNA status could explain the complete loss of SNV or its differential VAF in other samples. This filtering is essential to eliminate pseudo-heterogeneity being called among the multiple related samples. The second step is to cluster SNVs based on their CCF by using the Bayesian Dirichlet process-based clustering in a multidimensional mode (ndDPClust (https://github.com/Wedge-Oxford/dpclust); as previously described^5^) implemented based on DPClust v2.2.8 (https://github.com/Wedge-Oxford/dpclust) to identify clonal and subclonal clusters across multiple samples of the same patient. The DP clusters (identified as local peaks in the posterior mutation density) are then defined as clonal and subclonal according to their CCF peaks (with an expectation of one cluster at CCF of 1 representing clonal variants). Within individual samples, SNVs are annotated as clonal if they are assigned to the cluster with CCF of 1 and subclonal if assigned to a cluster with lower CCF. SNVs are annotated as truncal when they are clonal across all samples from a patient.

The third step is to construct phylogenetic trees across all samples by applying a set of rules to constrain possible trees^8,63^. Specifically, we assume that 1) the summed CCF of two clusters within a sample can only be above 1 if they occur in a co-linear fashion (i.e. on the same branch) while if below or equal to 1 may indicate co-linearity or branching, 2) clusters observed clonally within mutually exclusive sets of samples have a branching relationship (this is a special case of the ‘crossing rule’^63^) 3) in the presence of multiple clonal clusters unique to a subset of samples (i.e. branching), subclonal clusters can-not occur across all samples or in samples with branching lineages and are likely to be artefacts, and 3) within a branch, the temporal order of clusters is based on the magnitude of CCF, higher CCF clusters appearing earlier.

Given that all metastatic samples are clonally related, only one phylogenetic tree is constructed for each tumour. Individual sample trees are subtrees of the overall phylogenetic tree, which include just those subclones observed within a single sample. One of the strengths of multi-region sampling is that it exerts a greater inferential restriction on possible phylogenies, since the above stated rules must be simultaneously obeyed across all samples from a patient.

In tree construction, the relative branch lengths were made proportional to the fraction of all SNVs assigned to a cluster. We reconstructed the phylogenetic trees for all WGS- and WES-based patients using clusters representing at least 1% of the clustered SNVs. In tree visualisation, the percentage of all mutations in a cluster determined the relative branch length.

### Analysis of ITH in the primary tumour

As we only had targeted sequencing data on the two FFPE-based primary samples, and not WGS, we could not confidently call CNAs in these samples. To estimate CCF, we used the non-silent truncal SNVs (N=652) restricted to those that were in regions of the genome that were diploid in all metastatic samples (N=144, 22.2%; closely matching the global proportion of SNVs in diploid regions in all metastatic samples i.e. 24.6%) and diploidy was assumed in the primary samples for those loci. The density distribution of VAF was obtained for both primary samples and the peak with the highest VAF was inferred as the clonal set of variants. Purity was then estimated as twice the VAF of the clonal peak. With inferred CNA status and estimated purity, CCF for each SNV was estimated and ndDPClust was run on both primary samples to detect ITH.

### Driver mutation analyses

Melanoma drivers were identified as the 20 defined genes with relevant biological evidence outlined in a recently published seminal study^2^. Based on ndDPClust results, driver mutations present in the truncal cluster (i.e. present clonally across all samples of a patient) were assigned as truncal and those present in a non-MRCA cluster which formed a branch were assigned to that branch.

### Data and software availability/supplementary data

The targeted, whole genome and Affymetrix raw sequencing data have been deposited at the European Genome-Phenome Archive (EGA) (https://www.ebi.ac.uk/ega/ at the European Bioinformatics Institute). Data on all somatic SNVs, indels, inversions and copy number calls for both the index WGS autopsy case and the multi-site WES cases have also been deposited at the EGA. All these data are accessible via study ID’s EGAS00001001348 & EGAS00001003531 (as summarised below).

| EGA_StudyID | EGA_DatasetID | Patients | Data |
| --- | --- | --- | --- |
| EGAS00001001348 | EGAD00001005072 | Index_autopsy_case | Tumour/normal BAM files for all WGS data |
| EGAS00001001348 | EGAD00001005483 | Index_autopsy_case | Caveman, pindel, battenberg and brass calls |
| EGAS00001001348 | EGAD00010001717 | Index_autopsy_case | Affymetrix gene expression data |
| EGAS00001003531 | EGAD00001005073 | Index_autopsy_case | Targeted pulldown validation in support of the WGS analysis and targeted pulldown of the primary tumour |
| EGAS00001001348 | EGAD00001005421 | MultiSite_WES_Patients 1-7 | Tumour/normal BAM files for all WES data |
| EGAS00001001348 | EGAD00001005487 | MultiSite_WES_Patients 1-7 | Caveman, pindel and sequenza calls for all WES data |

All patients’ clinical details and sample mapping details are available in the **Supplementary Table**.

## Acknowledgements

We would like to thank the MelResist team for facilitating this research and providing the ethical framework for post-mortem donations, including Emily Barker, Doreen Milne, Catherine Wilson and Myfanwy Nicholas. The Cambridge Brain Bank for facilitating all aspects of the autopsy, particularly Jenny Wilson. The Tissue Bank team at Addenbrooke’s Hospital for helping to extract, store and deliver the samples including Martin Bromwich, Emily Daniels, Elizabeth Cromwell and Beverley Haynes. We thank Yvette Hooks for sectioning the primary tumour and digitally scanning the H&E images. The Cancer, Ageing and Somatic Mutation Programme team at the Sanger Institute for facilitating QC and DNA sequencing, particularly Claire Hardy, Stephen Gamble and Elizabeth Anderson. Computation used the Oxford Biomedical Research Computing (BMRC) facility, a joint development between the Wellcome Centre for Human Genetics and the Big Data Institute supported by Health Data Research UK and the NIHR Oxford Biomedical Research Centre. The views expressed are those of the authors and not necessarily those of the NHS, the NIHR or the Department of Health. Finally, we would like to sincerely thank the patients involved in this study and particularly the index patient and their family. Their complete dedication to advancing melanoma research despite an unpredictable and aggressive disease course dominated every clinical encounter. Their altruism and dedication to help has inspired every aspect of this work and lays the foundation for further studies of this kind.

## Author contributions

R.R. collected the clinical and molecular data, analysed the data and wrote the paper; N.A.P. analysed the DNA sequencing data, undertook the phylogenetic analyses and co-wrote the paper; O.C. helped analyse the Affymetrix expression data and performed the Consensus^TME^ clustering. D.L., F.S. and F.A.G. analysed the imaging data. D.L. also helped format the immune cell IHC images in Fig. 5. M.T. performed the histopathological analyses for the WES multi-site metastases patients. L.M. performed the laser-capture microdissection for the index patients’ primary tumour. I.F. helped select H&E images for Fig.1. K.W. analysed the copy number profiles for WES multi-site cases. J.M.M.G. and M.L. performed the immune cell IHC on the autopsy index patient’s samples. L.R. ran the mutational signature analyses. M.M. provided supervision on the expression analyses and Consensus^TME^ clustering. K.A. performed the autopsy and histopathological assessments of the index patients’ samples. P.C. provided overall clinical supervision including in study set up, patient recruitment as well as critical review of the manuscript. D.C.W. provided overall study supervision, particularly relating to the phylogenetic analyses and contributed to all sections of the manuscript. D.J.A. provided overall study supervision and critical review of the manuscript. All authors approved the final version.

